# Species richness, geographical affinities, and activity patterns of mammals in premontane Andean forests of the Magdalena river basin of Colombia

**DOI:** 10.1101/2020.07.26.221994

**Authors:** Diego A. Torres, Abel Eduardo Rojas

## Abstract

The Magdalena river basin is home to more than half of Colombia’s human population, and consequently the basin also harbors their economic activities. These activities have generated high deforestation rates and negative pressures on natural resources. With such a scenario of forest loss it is imperative to assess the state of the biodiversity and its conservation. Here, during six years we assessed the mammalian species richness and abundance in premontane forests of Caldas department in the Magdalena river basin. We also presented additional information on the activity patterns and geographical affinities of this fauna. We recorded 100 species of mammals with the Chiroptera as the richest order, followed by Rodentia. Most of the species are common and are not under risk of extinction; however, it is important to highlight the presence of six endemic species, three vulnerable species and one endangered species (white-footed tamarin). The mammalian fauna of this region is similar to other lowland localities in the Neotropics, and less similar to highland localities, including the nearby ones. Specifically, this fauna is most similar to lowland Tolima, and the Caribbean region of Colombia, Venezuela and Costa Rica; however, when we accounted only for bat fauna, it was more similar to the Caribbean and Pacific regions of Colombia. To secure the long-term persistence of these species we recommend maintenance of the current corridors such as riparian forests and living fences and an increase in the forested area.

## Introduction

The Magdalena river basin is located in northern South America in Colombia. This river has its origen in the Andes mountains (Colombian Massif) and flows northward between the Central and Eastern Andes in what is known as the Magdalena river valley, emptying finally into the Caribbean Sea. The geographic location in northern South America makes the biological diversity of this basin a combination of biotic elements with their origins from the main biographic regions of northern South America: the Andes, Amazon, Guiana Shield, Orinoquía and Caribbean lowlands (Velazco and Patterson 2013, Morrone 2014). As a warmer inter-Andean valley, the fauna of this basin has been associated with other lowland faunas with which it has a direct connection, such as that of the Chocó and Caribbean (Camacho et al. 1992, Kattan et al. 2004).

This basin is the most important area of the country, where much of the Colombian population and their economic activities are located (IDEAM 2001). Such a level of economic development has produced a continuous pressure on natural resources, resulting in high rates of deforestation mainly at low to mid elevations (Armenteras et al. 2013). It is estimated that around 70% of Andean and 30% of lowland forests in Colombia have been lost (Etter et al. 2006a, 2006b), much of the loss has been from the Magdalena river basin (Etter and van Wyngaarden 2000). This socio-ecological dynamic has created heterogeneous landscapes, where forests are represented mainly by fragments immersed in an agricultural matrix with different levels of connection through riparian forests and living fences (Etter et al. 2008). The remaining extensive areas of forests are mainly associated with the Andean highlands, national natural parks and private reserves associated with civil society as well as with hydroelectric dams (Armenteras et al. 2003).

As one of the mega-diverse countries of the world, it is of paramount importance to know the status of biodiversity conservation in such a scenario of Colombian forest loss. Important tools for this purpose are inventories and species monitoring resulting in checklists, natural history field data, and spatiotemporal trends in species richness and abundance. These data allow the construction and improvement of species distributional hypotheses (Elith et al. 2006), and also allows the evidence-based identification of important areas for conservation (Niemelä 2000). Moreover, checklists serve as historical data for tracking changes in future biodiversity studies.

Mammals are a very ubiquitous component of biodiversity in most ecosystems. They have a profound impact on the provision of ecosystem services due to a diversity of life histories (Ceballos and Ehrlich 2009, Kunz et al. 2011). In addition, many mammals are charismatic to people, and for this reason they are regularly used as umbrella or flagship species in conservation programs (Ducarme et al. 2013, Colléony et al. 2017). For these reasons biodiversity inventories generally include mammals. In the Magdalena river and its tributary rivers, mammal inventories can be tracked back as far as the 19th century (Mantilla-Meluk et al. 2014), and during the last decades checklists have been frequently published (e.g. Moreno-Bejarano and Álvarez-León 2003; Castaño and Corrales 2010; Garcés Retrepo et al. 2016; Solari et al. 2020). However, most published inventories and monitoring last less than one year and have low sampling efforts. Here, we provide a new checklist for mammals from premontane forests in the western slope of the Central Andes in the department of Caldas. Data were gathered for 6 years (2014-2019). To our knowledge, this is one of the most complete published inventories of mammals in the Magdalena river basin. We also provide some insights on the geographical affinities of this fauna as well as data on their activity patterns.

## Methods

### Study area

The forests included in this study are distributed on the eastern slope of the Central Andes in the department of Caldas, Colombia. We surveyed forests in the municipalities of Victoria, Norcasia and Samaná ranging in elevation between 300 and 1000 meters. This heterogeneous landscape is shaped from crops, pastures, and natural vegetation ranging from stubble to riparian forests and mature secondary forests. All sampling sites were located in the basin of the rivers Manso, Miel and Guarinó. These last two are tributaries of the Magdalena river. Sampling sites were under the influence of the area around the Miel I hydroelectric dam. The average temperature was 23°C with a maximum of 33°C with warmer conditions at lower elevations. Annual average precipitation varies from year to year between 3000 to 5000 mm, and distributed in an annual bimodal pattern with December to February and June to August as the dry periods (IDEAM 2001).

### Mammal sampling

We accumulated 647 days of sampling during the six years (2014-2019). The samplings where distributed during both rainy and dry seasons. For capturing bats we used mist-nets installed in the understory, across streams and at forest edges. Mist-nets remained open after sunset until 22:00 h. Manual captures were opportunistic, mainly associated with species roosting under small bridges, in hollow trunks, or on the foliage. For mist-nets a total of 34000 m-net/night was accumulated. For small and medium nonvolant mammals sampling we used live capture traps (Sherman and Tomahawk ®), located on the ground and up to two meters above the ground on branches. Traps were baited with a mixture of banana and oat, flavored with vanilla essence or sardine and corn with bacon butter. Each trap was verified daily in the morning and the bait replaced with fresh bait. The traps were installed in linear transects by stations 10 - 15 m apart, for a total of 20 - 25 stations according to the available area at each sampling site. Each station contained a trap on the ground and a trap in the branches. In total we accumulated 33455 trap/nights. Mammals were also sampled using trap-cameras (Bushnell ®) located along trials and streams where the passage of mammals was highly probable. Cameras were set 300 m apart at a minimum, and we accumulated a total of 3435 hours/cam. Direct observations were also included.

For bat nomenclature we followed Simmons and Cirranello (2020). For taxonomic updates in *Chiroderma* and *Tonatia* we followed Lim et al. (2020) and Basantes et al. (2020). The nomenclature of nonvolant mammals followed the Mammal Taxonomy Database of the American Society of Mammalogists (Burgin et al. 2018). For squirrels we followed the taxonomic arrangement proposed in Fiedler et al. (2020). Some specimens were collected, prepared as skull and skin, and deposited in the Museo de Historia Natural of the Universidad de Caldas. All procedures followed the guidelines of the American Society of Mammalogists for the use of wild mammals in research (Sikes and Gannon 2011).

### Data analysis

To establish geographical similarities of the mammalian fauna in the study area with other areas of the Neotropics, we constructed a matrix of presence/absence of 329 species of mammals from data available from checklists for localities in Central America and northern South America. We excluded the genera *Mazama* and *Sylvilagus* because it was not clear to what species to assign the species reported in the references. Specifically, we obtained data for the following areas: (1) the Central Andes of Colombia on the western slope in Risaralda (Castaño et al. 2018), Valle del Cauca (Rojas-díaz et al. 2012), Cauca (Ramírez-Chaves and Pérez 2010) and the eastern slope in Tolima (García-Herera et al. 2019); (2) for the Western Andes of Colombia on both slopes of Valle del Cauca and Cauca, and the Pacific lowlands (Ramírez-Chaves and Pérez 2010, Rojas-díaz et al. 2012); (3) for the eastern slope of the Colombian Massif in Cauca (Ramírez-Chaves and Pérez 2010); (4) for the Caribbean region of Colombia in Cordoba (Racero-Casarrubia et al. 2015); (5) for the Orinoquía of Colombia in Arauca (Mosquera-Guerra et al. 2019); (6) for the Guiana Shield in Colombia (Trujillo et al. 2018); and (7) for French Guiana (Lim 2012); (8) for the Amazonas in Venezuela (Lim 2012);(9) for the Sierra de Aroa in Yarucay State in northern Venezuela (García et al. 2016); (10) for Central America in Costa Rica (Rodríguez-Herrera et al. 2014); and (11) for La Rioja in Argentina (Fariñas-Torres et al. 2018); this last locality was chosen to be used as outgroup. We separated the mammal fauna in the localities of the Andes below 1000 m from the fauna above 2000 m of elevation. Localities were clustered based on the similarity index of Jaccard using the algorithm Paired Group (UPGMA) in the software PAST (Hammer et al. 2001).

We assessed inventory completeness as RO/RE*100, where RO was the observed species richness and RE was the species richness estimated by the index Chao 1 calculated with the software ESTIMATES based on a matrix of presence or absence of species and randomized 100 times (Colwell and Elsensohn 2014) using days as sampling units. As we did not intend to assess habitat use but only activity patterns, we used raw total abundances obtained in camera traps, constructing a frequency distribution graph for each of the 24 hours of the day for species with 20 or more records.

## Results

### Mammal richness

We gathered 9848 records of mammals representing 100 species from nine orders and 26 families. Chiroptera was the richest order with 53 species, followed by Rodentia with 19 species. Two species are listed globally as Endangered and Vulnerable (*Saguinus leucopus* and *Aotus griseimembra*, respectively), and four species are listed nationally as Vulnerable (Table 1; Appendix 1, 2, 3, 4). Six species are endemic to Colombia, most of them rodents (four spp.). The completeness of the inventory was 91.5% (Fig. 1). We expected to find according to the estimator Chao 1 at least nine species more if the sampling effort were increased. We presented a checklist for the area (Table 1), including two additional species (*Mustela frenata* and *Centronycteris centralis*) recorded for the area by Castaño and Corrales (2007 and 2010) but not recorded in this study.

**Table 1.**
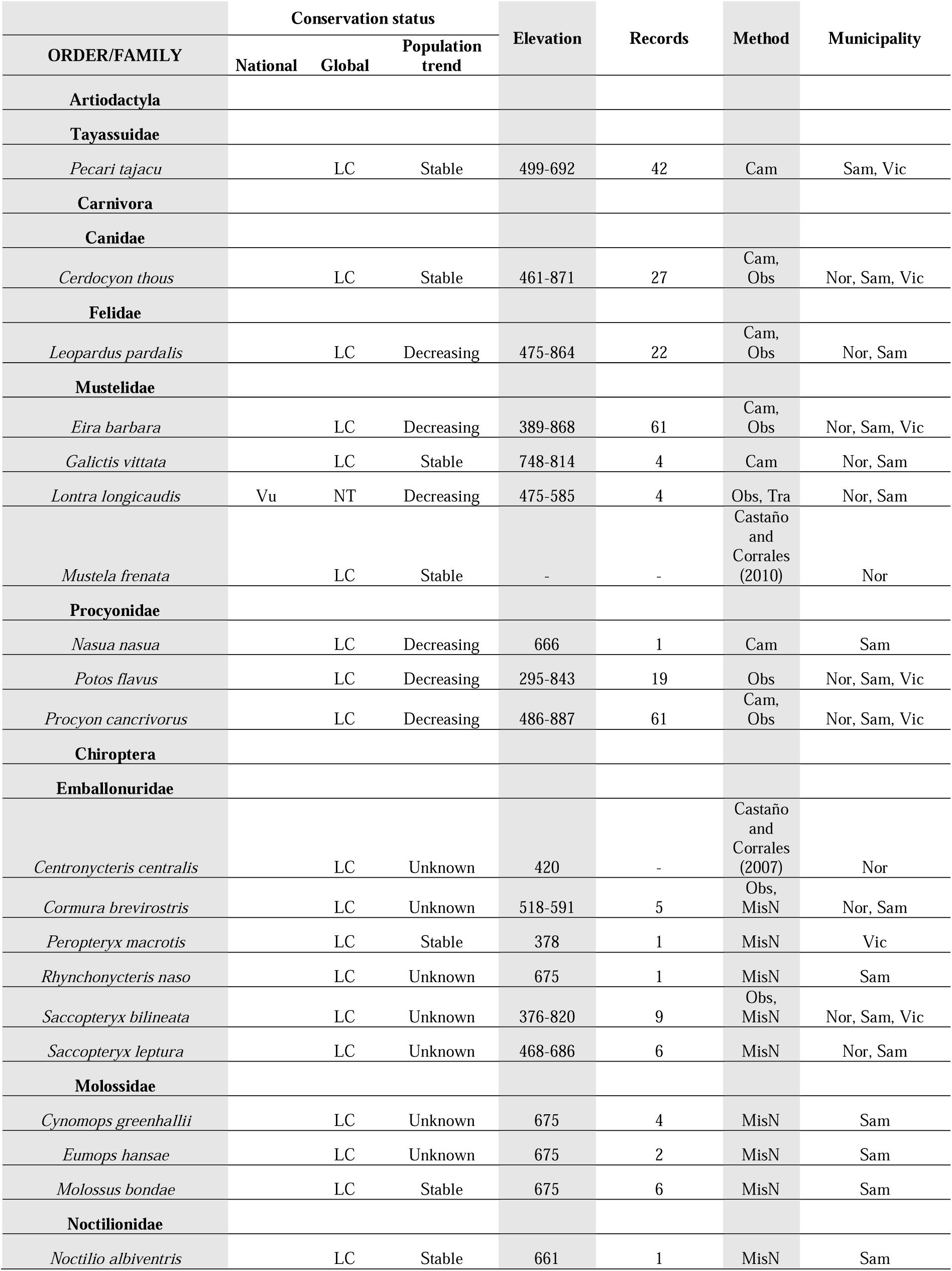
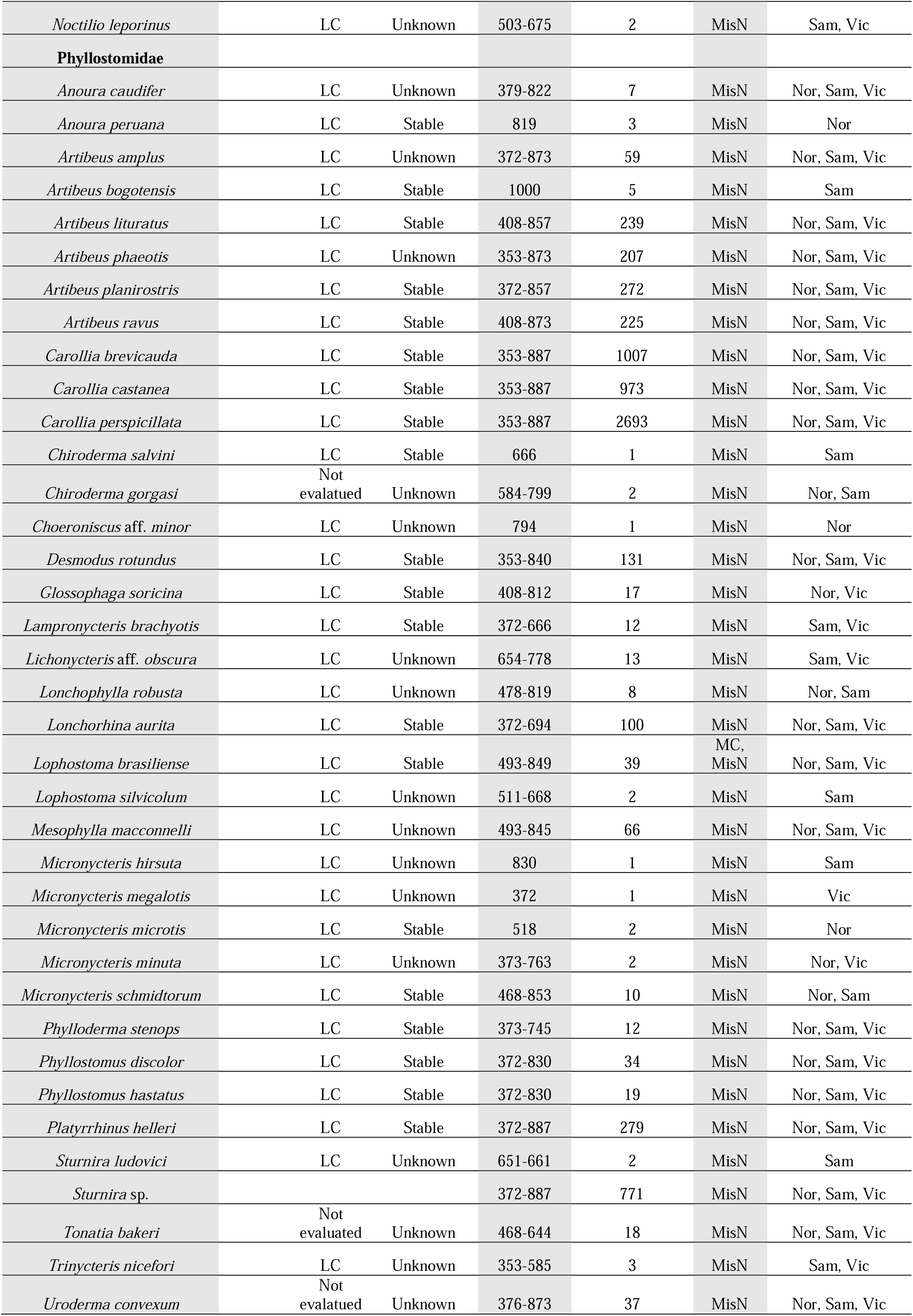
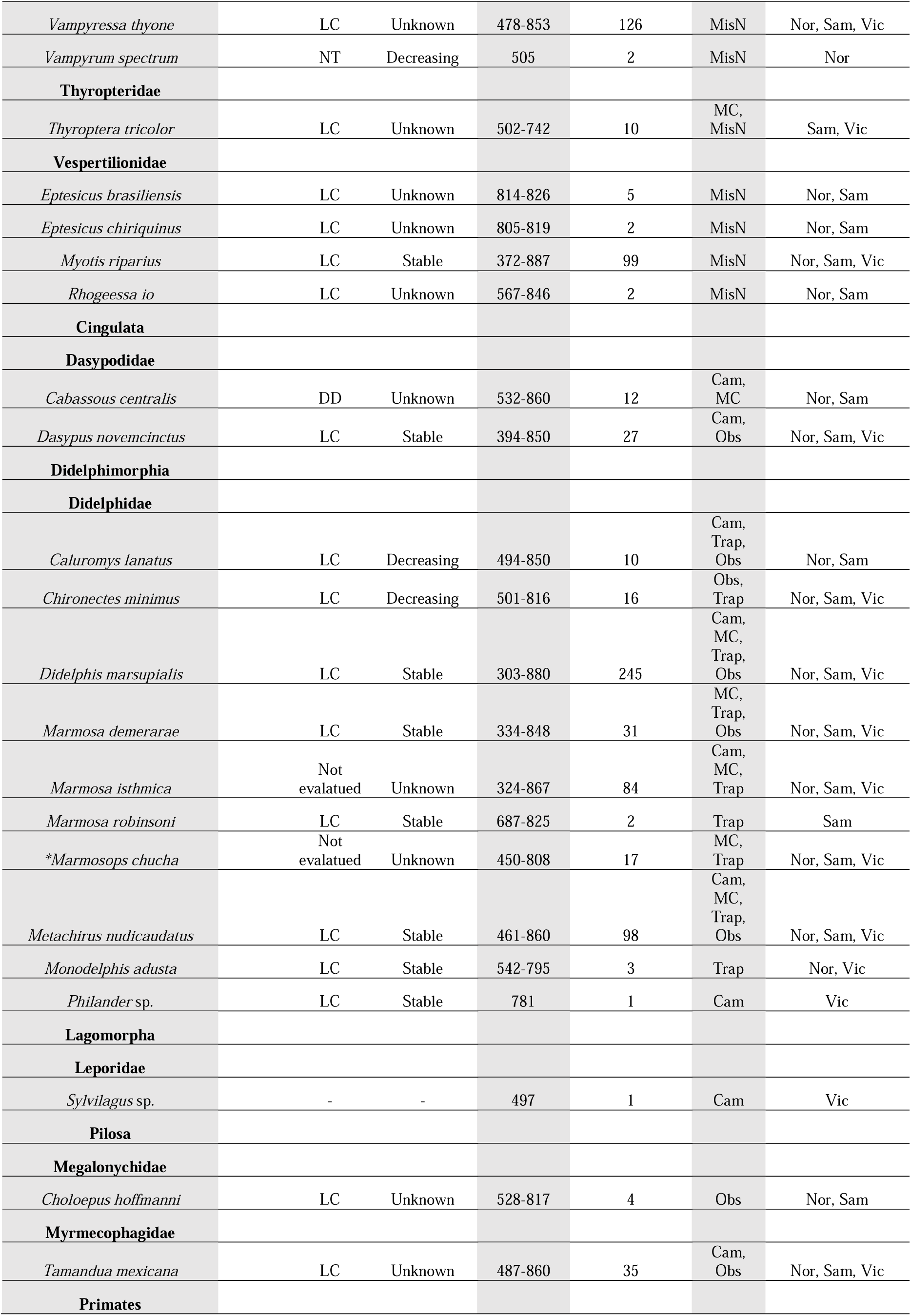
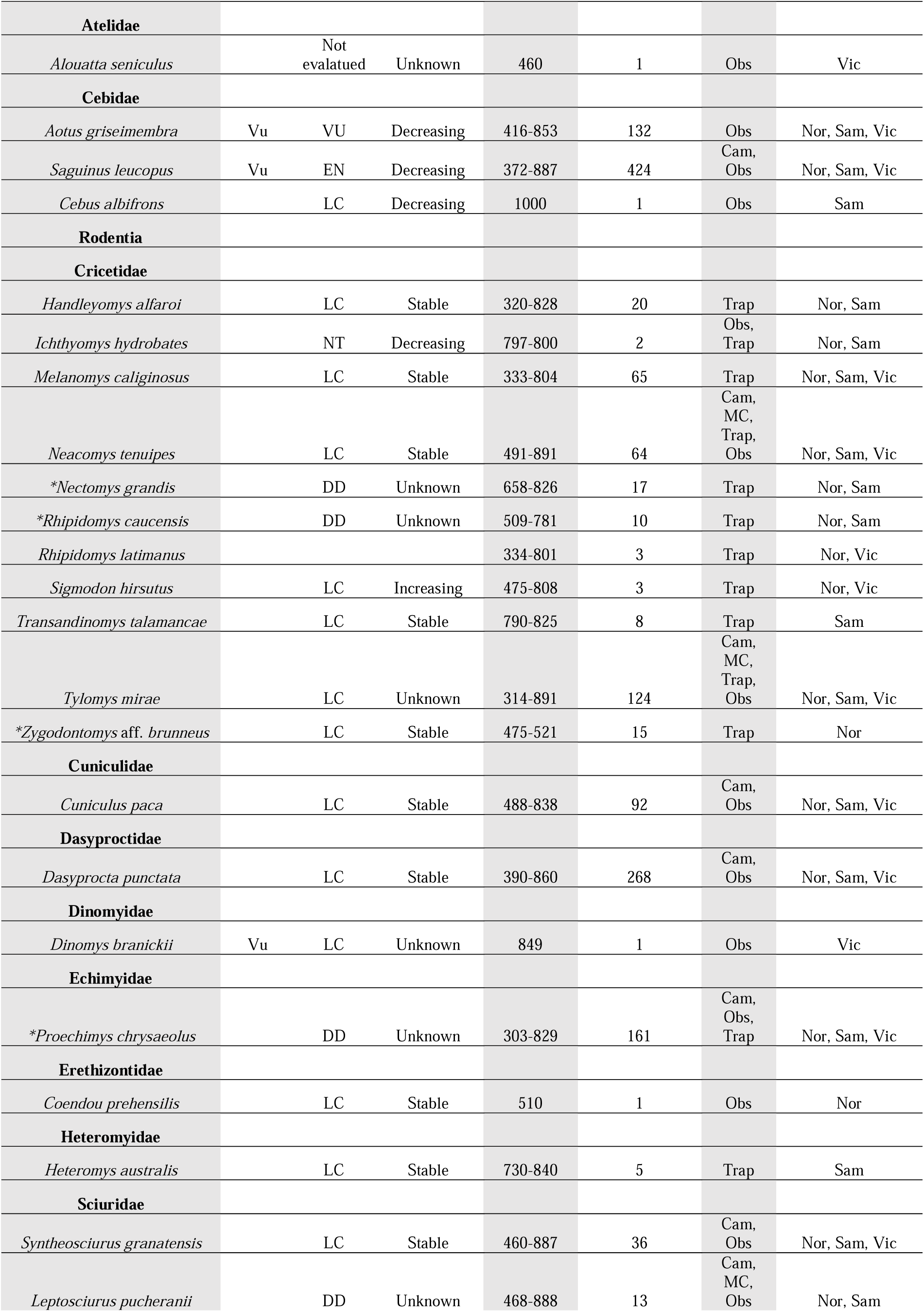
Checklist of mammals in premontane forests of the Magdalena river basin in eastern Caldas, Colombia. National (resolución 1912 de 2017) and global (Red List of IUCN) conservation status: **DD** (data deficient), **LC** (least concern), **NT** (near threatened), **VU** (vulnerable), **EN** (endangered). Methods: **Cam** (camera trap), **MC** (manual capture), **MisN** (mist net), **Obs** (direct observation), **Trap** (Sherman and Tomahawk traps). Municipalities: **Nor** (Norcasia), **Sam** (Samaná), **Vic** (Victoria).

**Figure 1.**
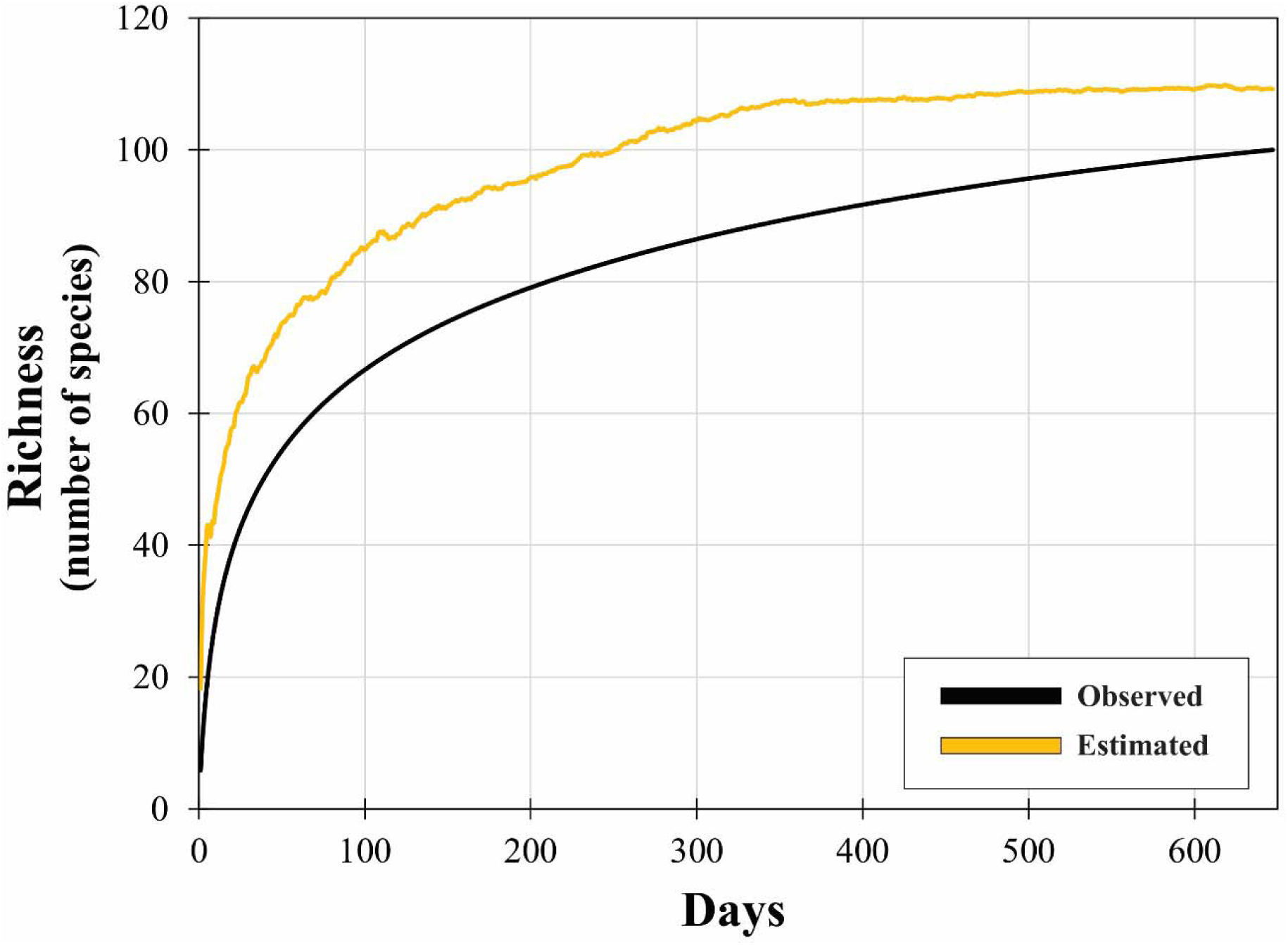
Curves of estimated (Chao 1) and observed species richness.

### Geographical affinities

The mammal fauna in the study area was most similar to the fauna of the same Central Andes eastern slope in Tolima below 1000 meters of elevation (Fig. 2A). Concurrently, the fauna of these two areas was similar to the fauna of the Caribbean region in Cordoba. Together, the fauna of these three regions was similar to that of the Sierra de Aroa in Venezuela and Central America in Costa Rica and formed a group with the fauna of the Venezuelan Amazon and Guiana Shield in Colombia and French Guiana. Altogether, this group was similar to another group containing the faunas of Central and West Andes below 1000 meters and above 2000 meters (Fig. 2A).

**Figure 2.**
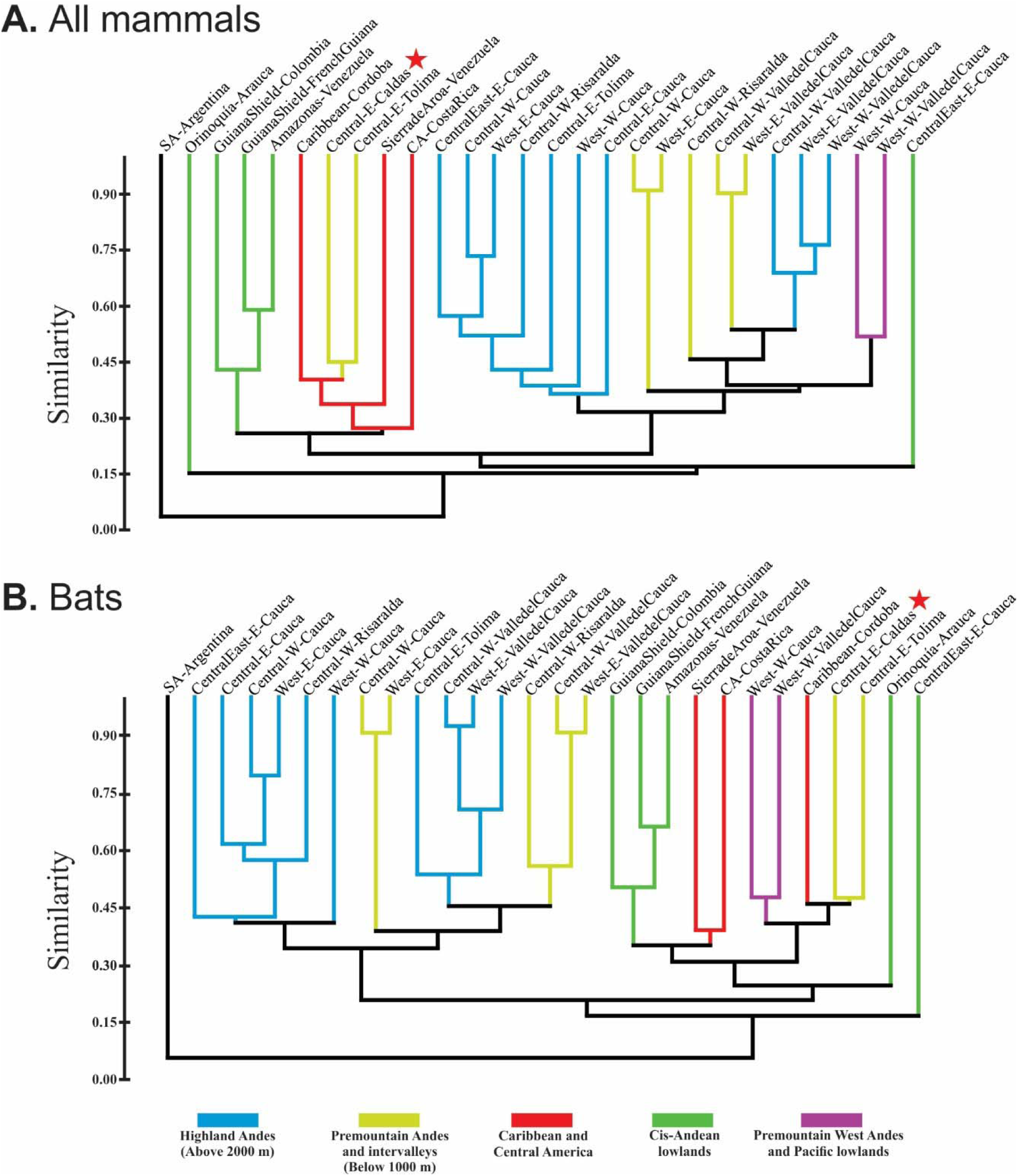
Dendograms showing the similarity in species composition (Jacckard index) of mammals (A) and bats (B) among northern South America and Central America localities. Red start indicate the study area. Central, West and East refers to the three ramifications of the Andes in Colombia; W (west) and E (east) refers to the slope of the Andes; SA (South America); CA (Caribbean); CentralEast refers to the Colombian Massif.

When only bat richness is considered, the pattern of similarity changes (Fig. 2B). Bat fauna was also more similar to the fauna of Tolima below 1000 meters, and these two regions were similar to the bat fauna of the Caribbean in Cordoba. However, instead of grouping with Central America or the Sierra de Aroa, the bat fauna of these three regions were more similar to bat faunas in the Pacific region of Valle del Cauca and Cauca. All regions making up this group were similar to other groups formed by the bat faunas of Venezuelan Amazon and Guiana Shield (Fig. 2B).

### Activity patterns

Mammals showed (Fig. 3) three types of activity pattern: (1) diurnal such as *Dasyprocta punctata* and *Eira barbara*, (2) cathemeral (active both at day and night) such as *Procyon cancrivorus*; and (3) nocturnal, like most species. However, some species showed activities that fell out of their regular activity pattern; for example, *E. barbara* showed activity in the first hours after dawn, and *Tamandua mexicana* and *Proechimys chrysaeolus* showed some activity during the day. All species showed some crepuscular activity.

**Figure 3.**
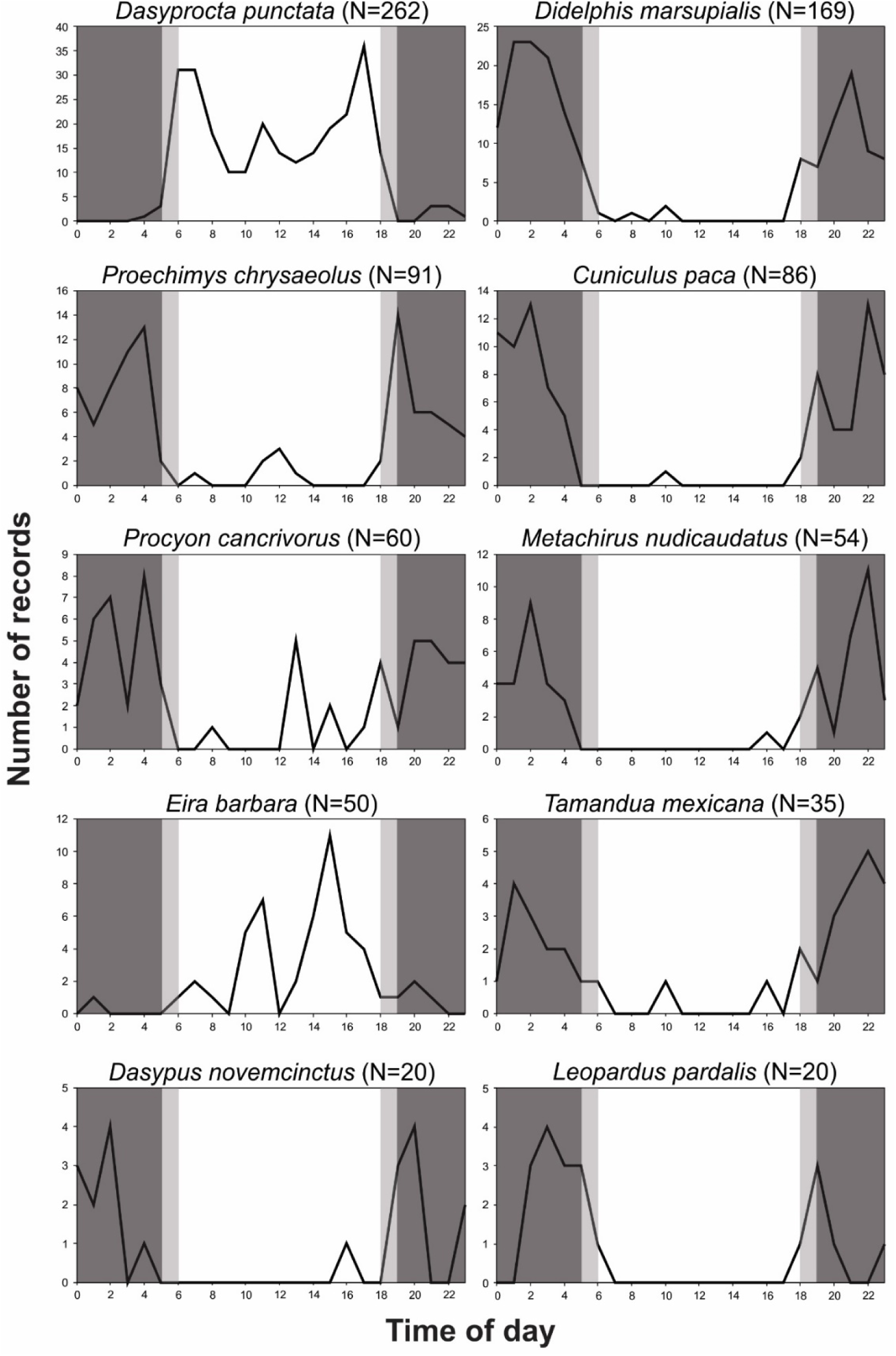
Activity patterns of some mammals in the study area. Dark gray areas indicate hours of darkness, while light gray indicate twilight. Records represent the observations in trap-cameras during the six years of monitoring.

## Discussion

Premontane forests located on the eastern slope of the Central Andes below 1000 meters of elevation in the department of Caldas sustain at least 100 species of mammals. This number represented 19% of the mammalian richness in Colombia (524 species, Sociedad Colombiana de Mastozoología 2017). Bats as the richest order and rodents as the second follow the same pattern found at the national scale (Solari et al. 2013). Based on the richness estimator Chao 1 is expected to find more species (nine species) if the sampling effort were increased. Mammal species richness checklists in localities in the Magdalena river basin report species that were not found in the study area, but their presence is highly probable, especially the animalivorous bats such as *Myotis caucensis, M. albescens, Pteronotus parnellii, Trachops cirrhosus*, some molossid species, and nectarivorous bats such as *Hsunycteris thomasi* and *Lionycteris spurrelli* (García-Herera et al. 2019, Solari et al. 2020). Some carnivorous mammals such as *Puma yagouaroundi* or *Bassaricyon neblina* are to be expected (Sánchez-Giraldo and Daza 2017, Gerstner et al. 2018). In addition to increasing the sampling effort, it is important to focus the effort on habitats where rare species concentrate their activities. This was the case of the bat *Eumops hansae*, which was captured over the river Manso in a net set across the river, a method that was rarely used in this or other studies; the specimen captured by this method was the first reported for the country (Torres and Rojas 2020).

In general, the mammalian fauna found in the study area is composed of common species with wide geographic ranges. However, the presence of six endemic small mammals with restricted geographic ranges associated with premontane and cloud forests in the Magdalena river basin (some including Cauca river) emphasize the conservation value of premontane forests in eastern Caldas department. Since nonvolant small mammals have low dispersal ability, local deforestation associated with agricultural activities can cause local extinctions; thus, a pattern is generated at the landscape scale of populations restricted to protected but generally isolated forested areas (Castro and Fernandez 2004, Paise et al. 2020). This is especially true for the marsupial *Marmosops chucha*, the rodent *Rhipidomys caucensis* and the primate *Saguinus leucopus* that carry out their activities above the ground in the lower, middle and upper strata of forests. To ensure the survival in the long term for these endemic species it is imperative to increase the connectivity of forest fragments and maintain the already existing corridors such as riparian forests and living fences (Zimbres et al. 2017).

Most species in the study area do not face a current risk of extinction (LC or NT), however, 10 species showed a globally decreasing population trend. This pattern is part of a world phenomenon known as defaunation, which means the progressive loss of animal populations caused mainly by habitat loss (Dirzo et al. 2014). It is interestingly to note that eight of the ten species with decreasing populations feed on other animals (i.e. animalivorous), despite belonging to different taxonomic groups such as bats (*Vampyrum spectrum*), carnivores (e.g. *Leopardus pardalis* or *Lontra longicauda*), marsupials (*Chironectes minimus*), and rodents (*Ichthyomys hydrobates*). Animalivory is a functional trait of species that have been associated with increased vulnerability to habitat loss and fragmentation because of their lower population densities and slow life histories (Cardillo et al. 2004, Minin et al. 2016).

As revealed by the faunistic similarity analysis, the mammalian fauna of the Magdalena river basin in eastern Caldas and Tolima below 1000 meters of elevation is related to the mammal fauna of the Caribbean region in Northern South America and Central America. This was expected because the valley of the Magdalena river between Central and East Andes exhibits warmer conditions that allow the establishment of typical species of lowland ecosystems not adapted to the cold of the uplands Andes (Soriano 2000, Kattan et al. 2004). This hypothesis related to temperature is also supported by the fact that mammalian faunas in the western Central Andes below 1000 meters and the Caribbean region are more related to the warmer lowlands of the Amazon and Guiana Shield, than they are to closer regions like West and Central Andes above 2000 meters. Our analysis restricted to bats also points to this relationship and adds to the group of “lowland faunas” the bat fauna of the Pacific region below 1000 meters. Similar relationships between lowland areas have been described for birds (Kattan et al. 2004, Cadena et al. 2016).

For birds it has been hypothesized that relationships among lowland areas in northern South America is mediated by dispersal events (Cadena et al. 2016). One hypothesis is that this dispersion occurred throughout the north in the Caribbean lowlands (Haffer 1967a, 1967b). Another hypothesis proposes that this occurred through passes in the Andes such as the Táchira depression, Andalucía pass and Suaza-Pescado valleys (Cadena et al. 2016). Good examples of this dispersion in mammals of the Magdalena river basin is the bat *Sphaeronycteris toxophyllum* with mainly a cis-Andean distribution, but some records exist in the middle Magdalena basin (García-Herrera et al. 2018). Also, the bat *Artibeus planirostris* found in the Amazon and Orinoco basins and beyond South America also extends its populations into the Magdalena basin (Solari et al. 2013). Thus, we suggest that dispersal events into the Magdalena river basin occurs both through north warmer corridors and passes throughout the Andes.

Most mammal species are active during the night, beginning at dusk and finishing at sunrise. This is the typical mammalian pattern; and, as it is currently understood, this was the ancestral behavior of the placental mammalian ancestor (Gerkema et al. 2013). Thus, activity during the day may be considered a derived behavioral trait. In general, the activity patterns of many Neotropical mammals of medium and large size are well-known (e.g. Blake et al. 2012; Ramírez-Mejía and Sánchez 2016; Huck et al. 2017); however, the patterns of small mammals are understudied. Here, we provide for the first time data on the activity patterns of the endemic spiny rat *Proechimys chrysaeolus*. This rodent showed intense activity immediately after dawn, and another peak of activity just before sunrise and some activity was evident at mid-day. This was the same pattern found in other species of *Proechimys* in the Brazilian Amazon (Pratas-Santiago et al. 2016).

## Conclusion

We conclude that the premontane forests of the Magadalena river basin in eastern Caldas harbor a rich mammalian fauna, conformed mostly of common species of lowland origins. However, these forests are of high conservation value because they host at least six endemic species and four endangered or vulnerable species. To secure the long-term persistence of these species we recommend maintaining the current corridors such as riparian forests and living fences and increasing the forested area.

## Acknowledgments

This study was financed by ISAGEN and the Universidad de Caldas (agreement 33/45). We thank Beatriz Edilma Toro, John Harold Castaño and Thomas Defler for valuable comments to this manuscript, and Mary Luz Bedoya for administrative and logistical support.

---

**Appendix 1.**
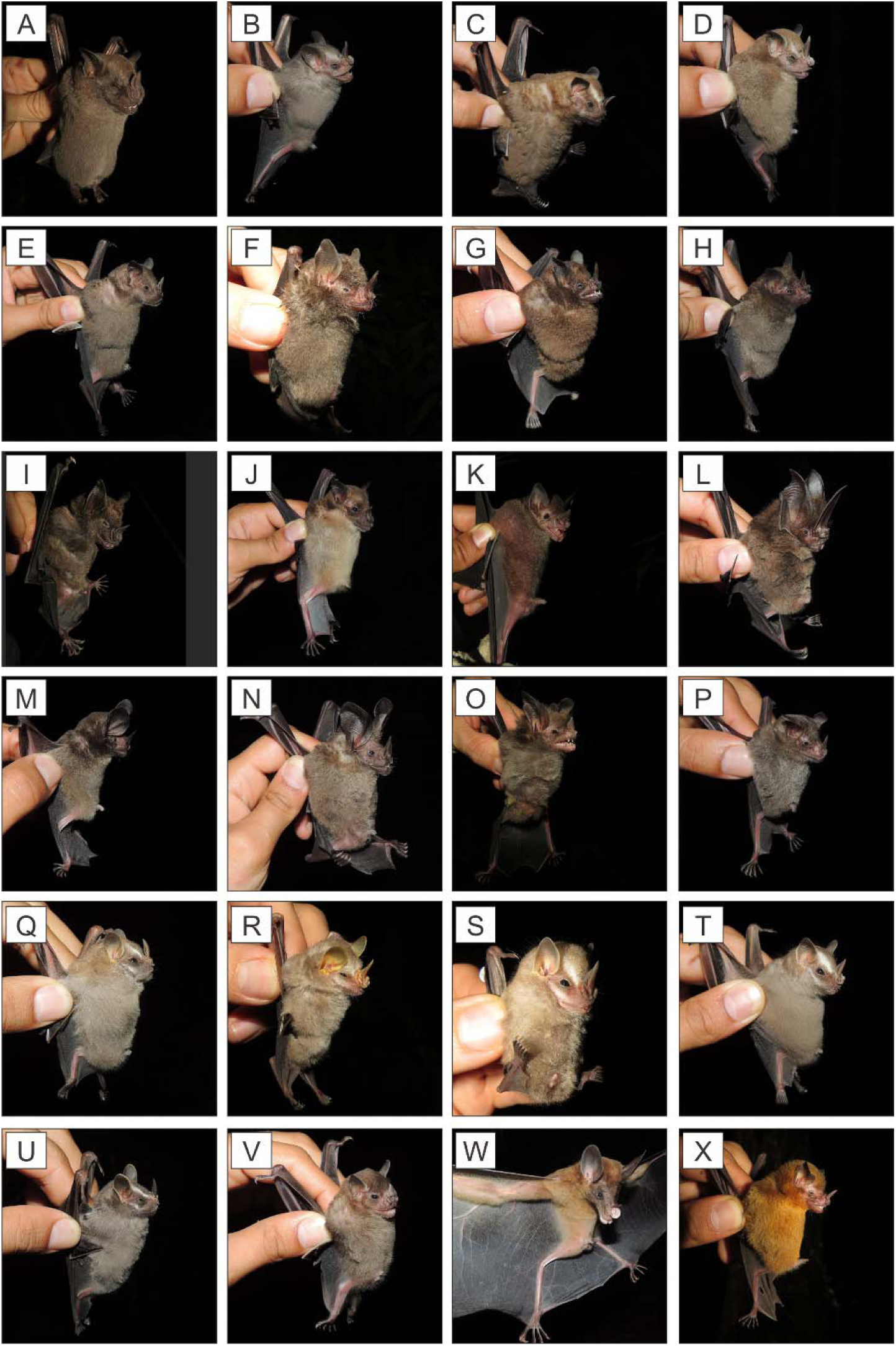
Some bat species captured in this study. *Artibeus amplus* (A), *A. phaeotis* (B), *A. lituratus* (C), *A. ravus* (D), *A. planirostris* (E), *Carollia brevicauda* (F), *C. castanea* (G), *C. perspicillata* (H), *Phyllostomaus hastatus* (I), *P. discolor* (J), *Phylloderma stenops* (K), *Lonchorhina aurita* (L), *Lophostoma brasiliense* (M), *L. silvicolum* (N), *Tonatia bakeri* (O), *Trinycteris nicefori* (P), *Chiroderma gorgasi* (Q), *Mesophylla macconnelli* (R), *Vampyressa thyone* (S), *Platyrrhinus helleri* (T), *Uroderma convexum* (U), S*turnira* sp. (V), *Vampyrum spectrum* (W), and *Lampronycteris brachyotis* (X).

**Appendix 2.**
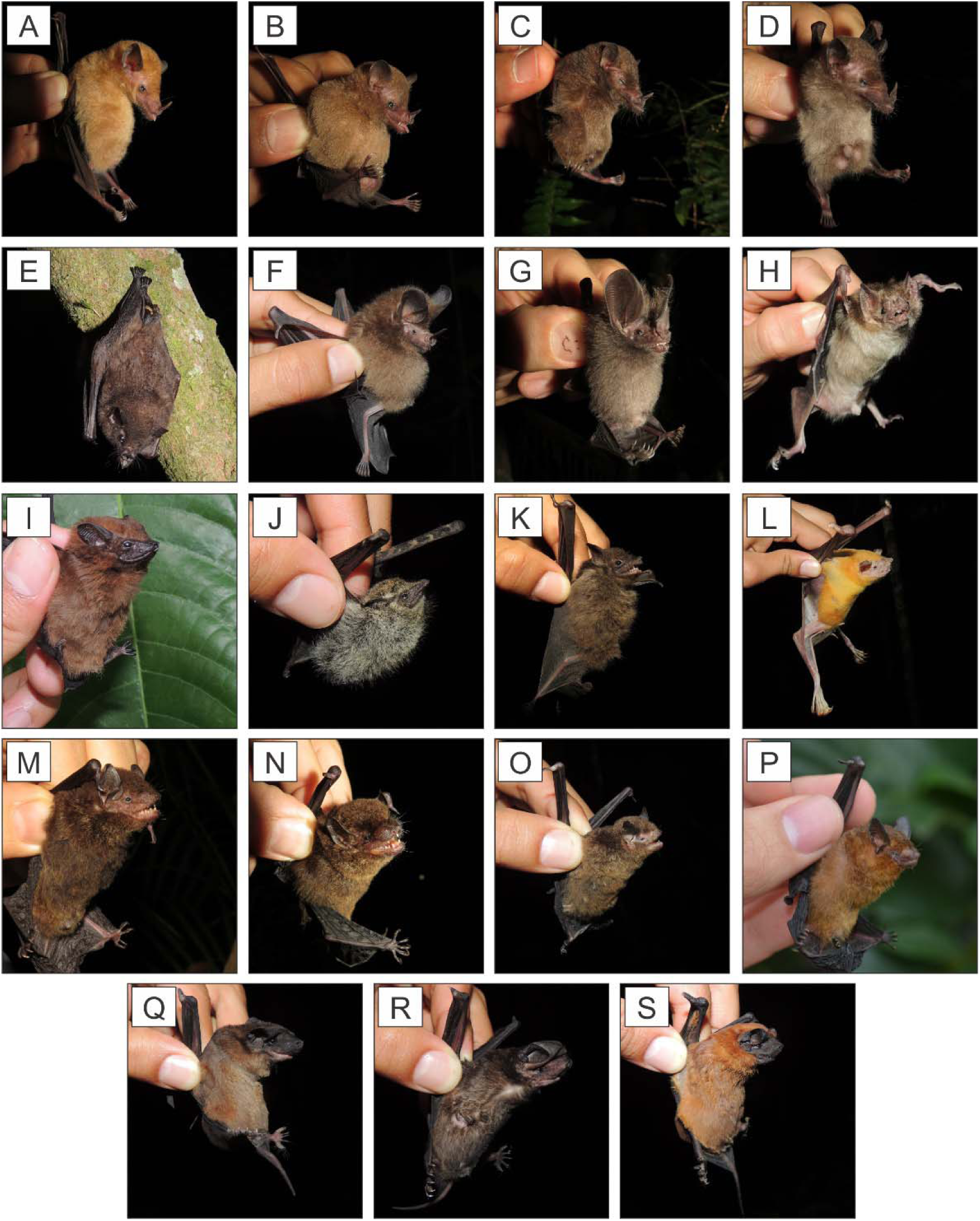
Some bat species captured in this study. *Lonchophylla robusta* (A), *Glossophaga soricina* (B), *Anoura caudifer* (C), *A. peruana* (D), *Lichonycteris* aff. *obscura* (E), *Micronycteris microtis* (F), *M. schmidtorum* (G), *Desmodus rotundus* (H), *Cormura brevirostris* (I), *Rhynchonycteris naso* (J), *Saccopteryx bilineata* (K), *Noctilio leporinus* (L), *Eptesicus chiriquinus* (M), *E. brasiliensis* (N), *Myotis riparius* (O), *Rhogeessa io* (P), *Cynomops greenhalli* (Q), *Eumops hansae* (R), and *Molossus bondae* (S).

**Appendix 3.**
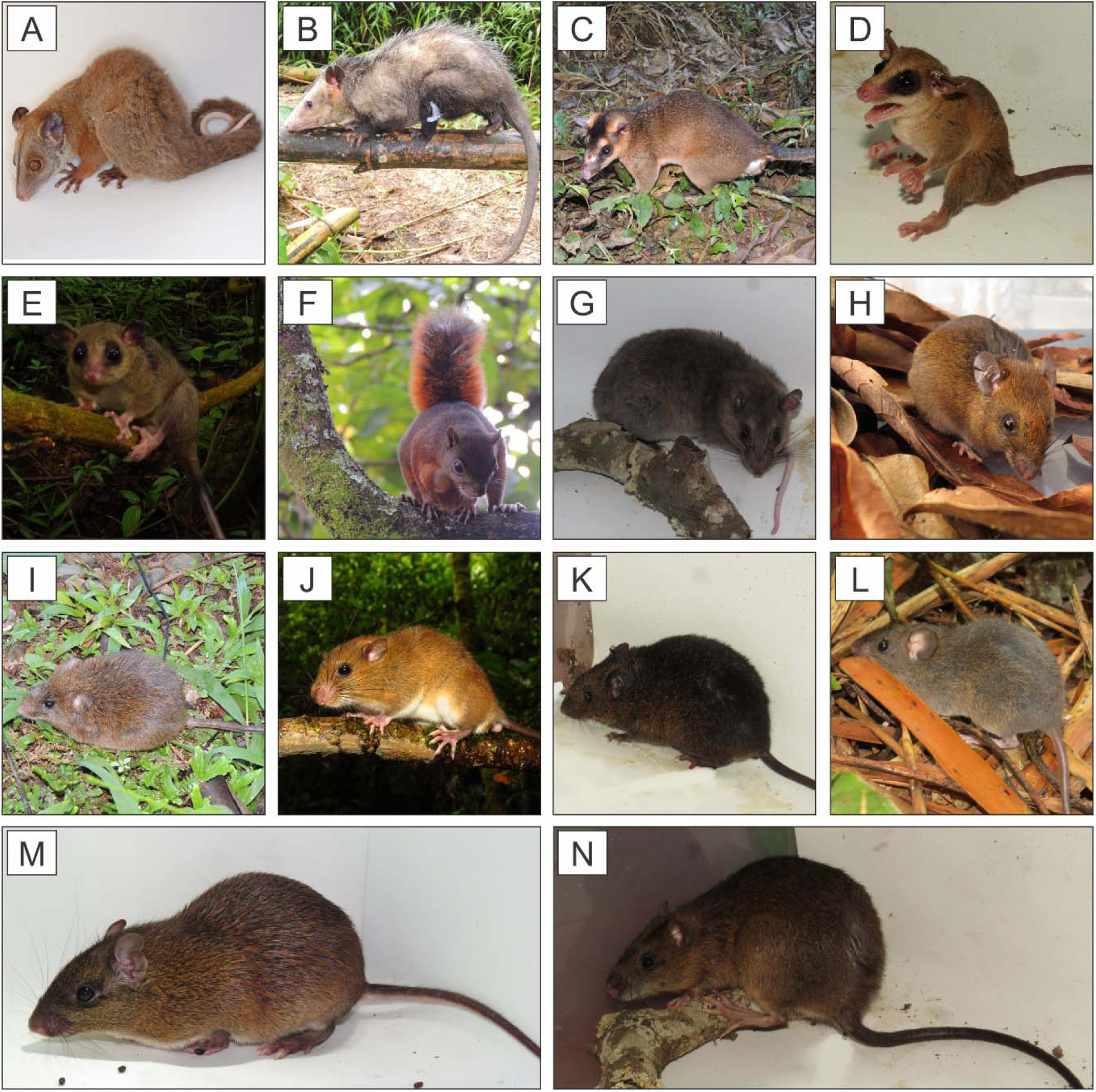
Some marsupials and rodents captured in this study. *Caluromys lanatus* (A), *Didelphis marsupialis* (B), *Metachirus nudicaudatus* (C), *Marmosa isthmica* (D), *M. demerarae* (E), *Syntheosciurus granatensis* (F), *Tylomys mirae* (G), *Transandinomys talamancae* (H), *Zygodontomys* aff. *brunneus* (I), *Rhipidomys caucensis* (J), *Melanomys caliginosus* (K), *Handleyomys alfaroi* (L), *Proechimys chrysaeolus* (M), and *Nectomys grandis* (N).

**Appendix 4.**
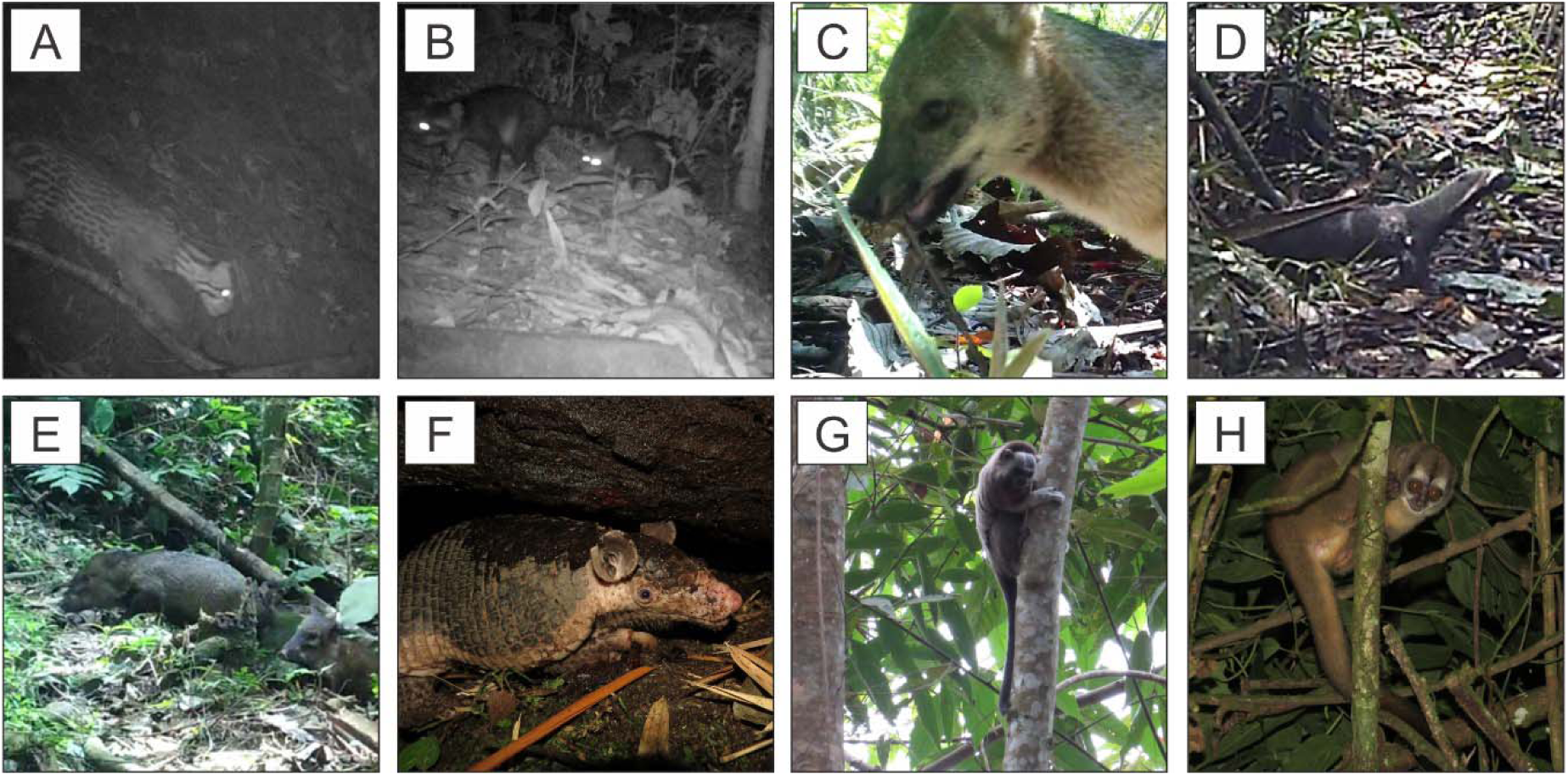
Some mammals registered in this study. *Leopardus pardalis* (A), *Procyon cancrivorus* (B), *Cerdocyon thous* (C), *Galictis vittata* (D), *Pecari tajacu* (E), *Cabassous centralis* (F), *Saguinus leucopus* (G), and *Aotus griseimembra* (H).

